# Microscopic characterization of local strain field in healing tissue in the central third defect of mouse patellar tendon at early-phase of healing

**DOI:** 10.1101/2021.05.26.445759

**Authors:** Eijiro Maeda, Kaname Kuronayagi, Takeo Matsumoto

## Abstract

Tendons exhibit a hierarchical collagen structure, wherein higher-level components, such as collagen fibres and fascicles, are elongated, slid, and rotated during macroscopic stretching. These mechanical behaviours of collagen fibres play important roles in stimulating tenocytes, imposing stretching, compression, and shear deformation. It was hypothesised that a lack of local fibre behaviours in healing tendon tissue may result in a limited application of mechanical stimuli to cells within the tissue, leading to incomplete recovery of tissue structure and functions in regenerated tendons. Therefore, the present study aimed to measure the microscopic strain field in the healing tendon tissue. A central third defect was created in the patellar tendon of mice, and the regenerated tissue in the defect was examined by tensile testing, collagen fibre analysis, and local strain measurement using confocal microscopy at 3 and 6 weeks after surgery. Healing tissue at 3 weeks exhibited a significantly lower strength and disorganised collagen fibre structure compared with the normal tendon. These characteristics at 6 weeks remained significantly different from those of the normal tendon. Moreover, the magnitude of local shear strain in the healing tissue under 4% tissue strain was significantly smaller than that in the normal tendon. Differences in the local strain field may be reflected in the cell nuclear shape and possibly the amount of mechanical stimuli applied to the cells during tendon deformation. Accordingly, restoration of a normal local mechanical environment in the healing tissue may be key to a better healing outcome of tendon injury.

## 1. Introduction

Tendon is a dense connective tissue that transfers muscle force to the bone during joint movement. Normal mature tendons mostly comprise type I collagen, with a small amount of type III collagen and other biomolecules, including other minor types of collagen, elastin, and other proteins and glycoproteins. Normal tendons exhibit a hierarchical structure of collagen from the molecular level to the level of fascicles, the basic functional unit of the tendon (Kastelic et al., 1978). This structural characteristic helps tendons deform without disruption by facilitating sliding between fascicles (Snedeker et al., 2009; Thorpe et al., 2015) and fibres (Cheng and Screen, 2007; Szczesny and Elliott, 2014; Thorpe et al., 2013). In addition, a limited amount of elongation of the fibres themselves and the recruitment and realignment of the fibres aid in deformation (Thorpe et al., 2013). Owing to the highly aligned structure of collagen fibres associated with these complicated behaviours, the nucleus of cells within the tendon matrix, tenocytes, acquires an elongated shape, and is subjected to the application of multimodal mechanical stimulations to tenocytes, including longitudinal stretch, transverse compression, physical shear, and interstitial fluid shear (Arnoczky et al., 2002; Lavagnino et al., 2008; Screen et al., 2003). In addition, tenocytes sense the presence and loss of tension imposed on collagen fibres, possibly by sensing the level of intracellular cytoskeletal tension reflecting the matrix tension level (Arnoczky et al., 2008; Lavagnino et al., 2003; Maeda et al., 2020, 2013b). These mechanical signals are prerequisite for tenocyte functioning to maintain tissue homeostasis.

It is well known that tendons possess only limited regeneration capability (Snedeker and Foolen, 2017). Once the tissue is injured, granulation tissue is regenerated at the repair site, which is followed by accumulation of collagen fibres and subsequent remodelling to tendon-like tissue. However, regenerated tissue does not exhibit the same tendon hierarchical structure and is thus considered a scar tissue. Scar tissue possesses a less well-structured network of collagen fibres and contains a greater amount of type III collagen than that in normal tendons (Dyment et al., 2013; Williams et al., 1980), with the absence of the hierarchical structure at the level of fascicles and above, an increased cell population with round nuclei, and low mechanical strength (Snedeker and Foolen, 2017). In a rabbit patellar tendon (PT) model, the tensile strength of the tissue regenerated in a central one-third defect of PT was only 18% of the strength of the normal PT at 3 weeks of healing, increasing to 57% at 24 weeks (Maeda et al., 2009). Previous studies with similar experimental models also resulted in the formation of scar-like tissue in tendon defects with insufficient mechanical strength (e.g. Dyment et al., 2012 used a mouse model and Awad et al., 2003 used a rabbit model). Approaches adopting the principles of tissue engineering and regenerative medicine have been pursued to achieve a satisfactory healing outcome in terms of recovery of the original tissue structure as well as mechanical properties. Although this research direction is promising, further investigation is warranted to understand why tendons cannot complete functional regeneration and end in scar formation. By understanding the scar healing mechanisms of tendons, it may be possible to control tendon healing, minimise scar formation and reaching a better healing.

Thus, the present study focused on the mechanical environment of cells in healing tendon tissue under physiological stretch, particularly the local strain field. As mentioned above, tenocytes in normal tendons are exposed to mechanical stimuli, which are essential for the onset of mechanotransduction events and the subsequent expression of genes and proteins for the maintenance of tissue homeostasis (Wang, 2006). Mechanical stress is also an essential factor in achieving better functional tendon healing. Regenerated tissue in a central defect of rabbit PT healed under a stress-deprived condition only gained strength at a level of 5% of the normal strength at 3 weeks. However, when mechanical stress was re-applied after 3 weeks, at 12 weeks, it acquired a strength similar to that under the normal loading condition (Maeda et al., 2009). This implies that cells in the healing tissue can also sense the presence/absence of mechanical loading and respond to the mechanical loading applied to the tissue. Therefore, this leads to the hypothesis that a lack of collagen fibre dynamics (elongation, shear, and rotation) in healing tendon tissue may result in a limited application of mechanical stimuli to cells in the healing tissue, leading to incomplete recovery of tissue structure and functions in regenerated tendons.

It has previously been demonstrated that the magnitude of cell nucleus deformation is significantly smaller in the tissue regenerated in tendon defects than in the normal tendon (Freedman et al., 2018), suggesting that the cells in the regenerated tissue are exposed to less mechanical stress. Nonetheless, the local strain field in tendons has only been well characterised in healthy, normal tendon tissues, and the details of tissue local strain have not been clarified in healing tendon tissue.

Therefore, the present study was performed to measure the microscopic strain field in healing tendon tissue to facilitate a better understanding of the mechanical environment of cells within the healing tissue.

## 2. Materials and methods

### 2.1 Animal model

In the present study, 10-week-old male mice (ddY strain, Japan SLC) were used as the experimental animals. All animal experiments were approved by the Institutional Review Board for Animal Care at the Graduate School of Engineering, Nagoya University (approval #18-8, #19-10, and #20-3) and were performed in accordance with the Guide for Animal Experimentation, Nagoya University.

Each animal was subjected to surgery for the creation of a one-third defect in the middle of the PT of the right knee, using a method similar to that used previously for a rabbit PT model (Maeda et al., 2009). Briefly, under anaesthesia and sterile conditions, the right PT was exposed through a skin incision. A blunt stainless-steel blade was inserted beneath the PT through small incisions made laterally on both sides of the PT, and an infrapatellar fat pad was separated from the posterior surface of the PT. A rectangular defect was created in the middle of the PT, with a length spanning from the attachment of the patella to that of the tibia and a width of approximately one-third of the whole PT width (approximately 0.5–0.6 mm). The incision was closed with a 5-0 nylon suture, followed by sterilisation of the skin wound with povidone-iodine. The left knee was not operated on and therefore served as a contralateral control. All operated animals were allowed unrestricted activities in cages for 3 or 6 weeks.

### 2.2 Bulk tissue mechanical properties

To determine mechanical properties, both healing tissue filling the defect and the contralateral PT tissue were subjected to tensile tests. At the end of the designated healing periods, animals were euthanized using carbon dioxide gas, and a patella–PT–tibia complex was collected from both limbs. Samples were stored at −20°C until biomechanical analysis. A specimen for the tensile test was prepared by removing all unnecessary soft tissues from the complex. The complex was then trimmed to prepare a specimen with a dumbbell shape using a biopsy punch. In case of the healing tendon, residual tendon tissues on both sides of the defect were trimmed. The cross-sectional area of the middle portion (gauge section) of the specimen was measured using a non-contact image-based technique, as previously described (Tohyama et al., 2003). The complex was rotated in a saline bath using a stepping motor, and the front view of the tendon tissue was captured with a CMOS camera (acA1920-40uc, Basler, USA) at 18° intervals, using programs written in Arduino and LabVIEW (National Instruments, USA). The width of the tendon in each image was measured using ImageJ, and the cross-section was reconstructed to calculate the area.

Tensile tests were performed using a well-established method in a rabbit PT model (Yamamoto et al., 1992). The tibia of the complex was embedded into an aluminium cup with polymethylmethacrylate resin, and the patella was bound by two acrylic plates fabricated with a 3D printer (Keyence, Japan) and designed to hold the patella. The complex was set onto a tensile testing machine equipped with a 20 N load cell (EzTest, Shimazu, Japan), using a metal chuck for the patella and a custom-made stainless-steel jig for the tibia cup. Two parallel lines were drawn transversely, 1-mm apart, on the surface of the gauge section in each specimen using nigrosine (Sigma-Aldrich, USA). These lines were used for strain measurements following the test.

During testing, the specimen was immersed in a physiological saline solution warmed to 37°C. Following the application of several loading cycles between 0 and 0.1 N to the specimen as preconditioning, the specimen was stretched until failure at a stretch rate of 10 mm/min (the lowest value was set in the testing machine). The load applied to the specimen and the front view of the gauge section were recorded using a CMOS camera and a custom-made LabVIEW. Stress was calculated from the tensile load divided by the initial cross-sectional area of the gauge section, and the strain was determined from the increase in the distance between the gauge length markers divided by the initial distance. The peak stress was defined as the tensile strength, and the tangent modulus was defined as the slope of the stress– strain curve between 20% and 60% of the tensile strength.

### 2.3 Two-photon laser microscopy for collagen fibre alignment analysis

A subgroup of operated animals was assigned for the analysis of collagen fibre structure in both normal and healing tendon tissues using two-photon microscopy. At the end of 3 and 6 weeks of the healing period, the patella–PT–tibia complex was prepared from both sides of the knee, as described above. The complex was fixed in 4% paraformaldehyde solution in phosphate-buffered saline (PBS; Wako, Japan) for 3 h at room temperature (20-25°C); the PT was straightened with a 1 g weight. Following cell nucleus staining with 4 µM ethidium homodimer-III (Nacalai Tesque, Japan) in PBS overnight, the complex was immersed in PBS containing 60% w/w iodixanol (Optiprep, Abbott Diagnostics Technologies AS, USA) at 4°C overnight for tissue clearing. The PT section (or healing tissue section) was visualised using a two-photon laser microscope (Nikon A1R MP, Nikon, Japan) and a 25× water immersion objective lens. The excitation wavelength was 860 nm, and the second harmonic generation (SHG) signal from collagen and the fluorescent signal from the cell nuclei were obtained with a photomultiplier tube fitted with a 492SP filter and 575/25 filter (both from Nikon), respectively. The images obtained were 512 × 512 pixels, corresponding to a 520 × 520-µm field of view. A z-stack with a 2-µm step from the tissue anterior surface to approximately 200-µm deep was obtained at four to six locations longitudinally covering the mid-portion of the PT or healing tissue.

Collagen fibre orientation was analysed as an indicator of tissue integrity using the Fiji command Dimensionality. A representative image slice of the z-stack was selected, and a small square region of interest (ROI), consisting of 64 × 64 pixels, was cropped every eight pixels along both the vertical and horizontal directions to make a sub-stack, resulting in an image stack of 3249 ROI images. The orientation angle of collagen fibres, visualised by SHG signals, in each ROI image was analysed with Directionality. This performed two-dimensional fast Fourier transform (2D-FFT), and the probability density function of the Gaussian distribution was fitted to the angular distribution of collagen fibre orientation. From the analysis, three parameters describing fibre orientation were obtained: the peak (degree) and dispersion (unitless) of the fitted probability distribution function as well as the goodness of fit (a unitless value ranging between 0 and 1). Each pixel of the ROI image was assigned a peak and a value describing goodness of fit of the image, and this information was then brought back to the pixels corresponding to the original 512 × 512 image. As most of the pixels in the original image were analysed with Directionality multiple times, the representative peak angle of the pixel was calculated as follows:

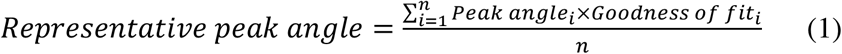

where *n* is the number of times analysed.

To avoid unnecessary errors in the analysis of the collagen fibre orientation, the pixels applicable to either of the following conditions were eliminated: 1) Directionality analysis was performed only once, and 2) the pixel intensity of the original image was not more than 50. The frequency distribution of the representative peak angle in the rest of the image pixels was fitted with the probability density function of the von Mises distribution using MATLAB (Mathworks, USA) to obtain the peak angle, *μ*, and a concentration factor, *κ*, of the distribution; which is analogous to the inverse of the variance of the Gaussian distribution, and thus, a high *κ* value indicates an angular distribution with less variability. We adopted this approach because it clearly shows the regions of the image with the peak angle of the fibre alignment, which are not accessible by performing a conventional 2D-FFT analysis of an entire image.

### 2.4 Confocal microscopy under tensile loading

Another subgroup of operated animals was assigned for confocal microscopy and local strain analysis. The patella–PT–tibia complex was obtained from both knee joints at the end of the designated healing period, and all soft tissues other than the PT were removed from the complex. The cell nuclei in the PT tissue were stained with Hoechst33342 (Invitrogen, USA), 1/1000 diluted in PBS, for 20 min at room temperature, followed by a brief wash in PBS. The patella and tibia were held by a custom-made acrylic grip for stretching on a confocal laser scanning microscope (IX81+FV1200, Olympus, Japan). The sample and grips were assembled in a silicone chamber (STB-CH-04, Strex, Japan), and the sample was kept hydrated with PBS within the chamber. A pair of cartridge heaters was inserted into the chamber to maintain the specimen at 37°C during imaging. The chamber, containing the specimen attached to the grips, was connected to a custom-made stretching device, equipped with a pair of linear actuators (Oriental Motor, Japan) operated using a custom-made LabVIEW program (Maeda et al., 2013a).

Before the stretching experiment, the specimen was preconditioned by repeatedly stretching to straighten and slightly stretch the collagen fibres and then unstretched to a relaxed state several times under the observation of fluorescence from the cell nuclei through the eyepiece with a 10× objective lens, and the specimen was stretched again to set the zero-strain point where the apparent slackness of the tendon structure was removed. In the deformation analysis, the cell nuclei within the tissue were visualised with a 20× objective lens (Olympus), with ×2.5 zoom and an excitation laser at a wavelength of 405 nm. A field of view (800 × 800 pixels, corresponding to 254.46 µm square) was selected in the middle of the tissue section as a representative region. The specimen was stretched to 700 µm (grip–grip distance) with an increment of 100 µm at a rate of 20 µm/s. At each stretching step, including the zero-strain point, a series of z-stacks was obtained from the tissue surface (z = 0 µm) to the inside of the tissue (z = approximately 50–100 µm, depending on the visibility of each sample), with an interval of 5 µm.

### 2.5 Image analysis for microscopic strain field

To characterise the microscopic strain field in both normal and healing tissues in the tendon, the local strain distribution was determined. A series of z-stack images obtained from each stretching step of confocal microscopy was projected onto a single image using maximum intensity projection, and a total of eight projected images were stacked. In the series of stacked images from 0 µm to 700 µm stretch, a set of ROIs were created for the cell nuclei that could be traced in all eight images in the series. The ROIs were used to calculate the xy-coordinates of the centroids of each cell nucleus using MATLAB.

The coordinates for the first frame (0-µm stretch) were used to construct triangular meshes using the MATLAB Delaunay function. Green-Lagrange strain tensor was calculated for each mesh at each stretching step, in comparison to the initial coordinates. Strains in the longitudinal and transverse directions (*E*_*xx*_ and *E*_*yy*_) as well as shear strain (*E*_*xy*_) were calculated using the following equation:

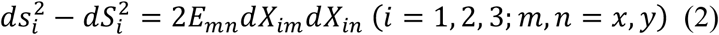

where *ds*_*i*_ and *dS*_*i*_ are the lengths of the *i*th side of a triangular element, respectively, and *dX*_*im*_is the x- or y-coordinate of the *i*th vertex of the triangle at the zero-strain position.

In addition, the microscopic tissue strain *E* was determined in the images from the changes in the distance between pairs of the cell nuclei positioned at almost identical vertical positions but located at a distance of approximately 200 µm along the horizontal direction (tissue long axis). Cell nuclear shape was evaluated in the long axis to the short axis ratio (nucleus aspect ratio, NAR) at the zero-strain level, and its change (*Δ*NAR) was evaluated at 4% microscopic tissue strain as NAR_*E*=4%_/NAR_*E*=0%_ −1.

### 2.6 Statistical analysis

Statistical analyses were performed using statistical language R (ver. 4.3.0). Tensile strength, tangent modulus, and strain at failure were examined using one-way analysis of variance (ANOVA), followed by Tukey’s honestly significant difference multiple comparisons if statistical significance was confirmed by one-way ANOVA. Green-Lagrange strain components, collagen fibre alignment parameters (peak and concentration), cell nucleus aspect ratio, and *Δ*NAR were examined using the nonparametric Kruskal–Wallis test, followed by Steel–Dwass multiple comparison test if statistical significance was found in Kruskal–Wallis test. In all tests, the significance level was set at *p* < 0.05.

## 3. Results

### 3.1 Bulk mechanical properties

Figure 1 shows the stress–strain relationships for the control group (CTL) and healing tissue at 3 weeks (H3w) and 6 weeks (H6w). The relationships for the healing tissues had a lower slope and breaking point than that for the normal tendon. The tensile strength and tangent modulus (mean ± standard deviation) of H3w (2.5 ± 1.4 and 38.7 ± 19.0 MPa, respectively) were significantly lower than those of CTL (17.0 ± 4.0 and 283.7 ± 176.1 MPa, respectively) (Fig. 2), and the values were still significantly lower than those of CTL at 6 weeks (3.8 ± 2.0 and 58.3 ± 36.4 MPa, respectively). There were no significant differences between the H3w and H6w groups (Fig. 2). There were also no statistically significant differences in strain at failure among the three groups (0.09 ± 0.02, 0.10 ± 0.02, and 0.10 ± 0.02 for CTL, H3w, and H6w, respectively).

**Figure 1.**
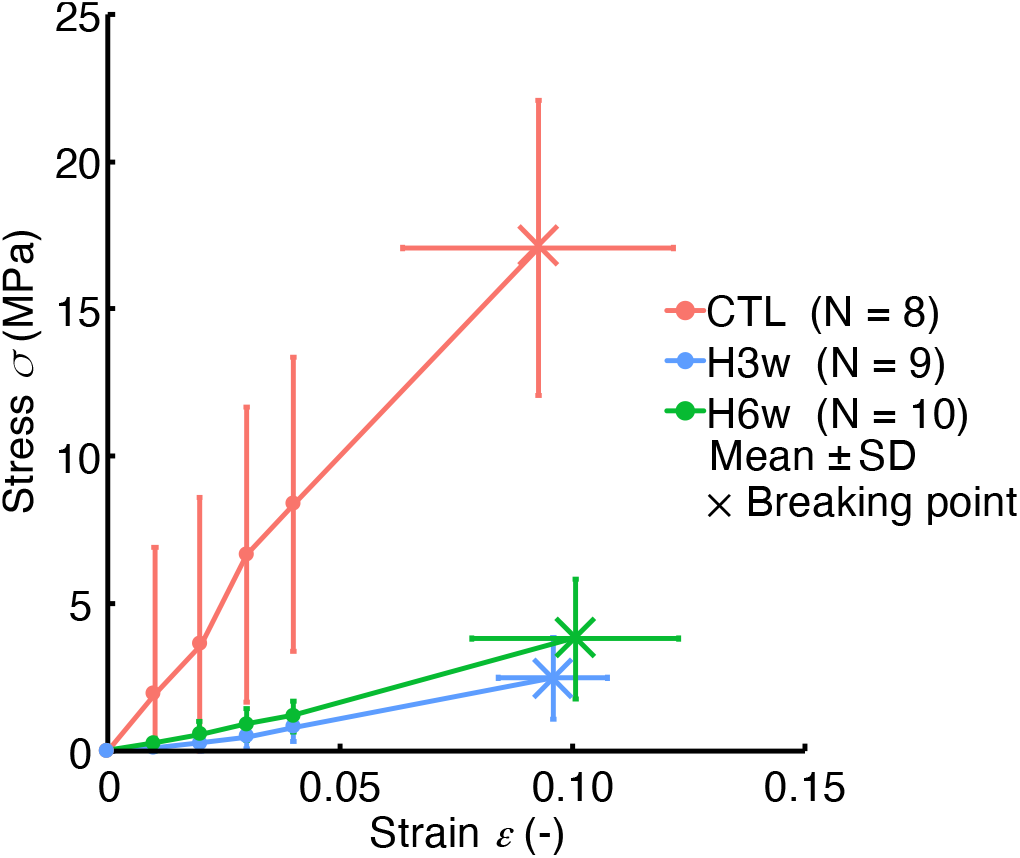
Stress–strain relationships of mouse patellar tendons and their healing tissues. CTL, contralateral control PT; H3w, healing tissue regenerated in the PT defect examined at 3 weeks; and H6w, healing tissue regenerated in the PT defect examined at 6 weeks; N, number of specimens; SD, Standard deviation.

**Figure 2.**
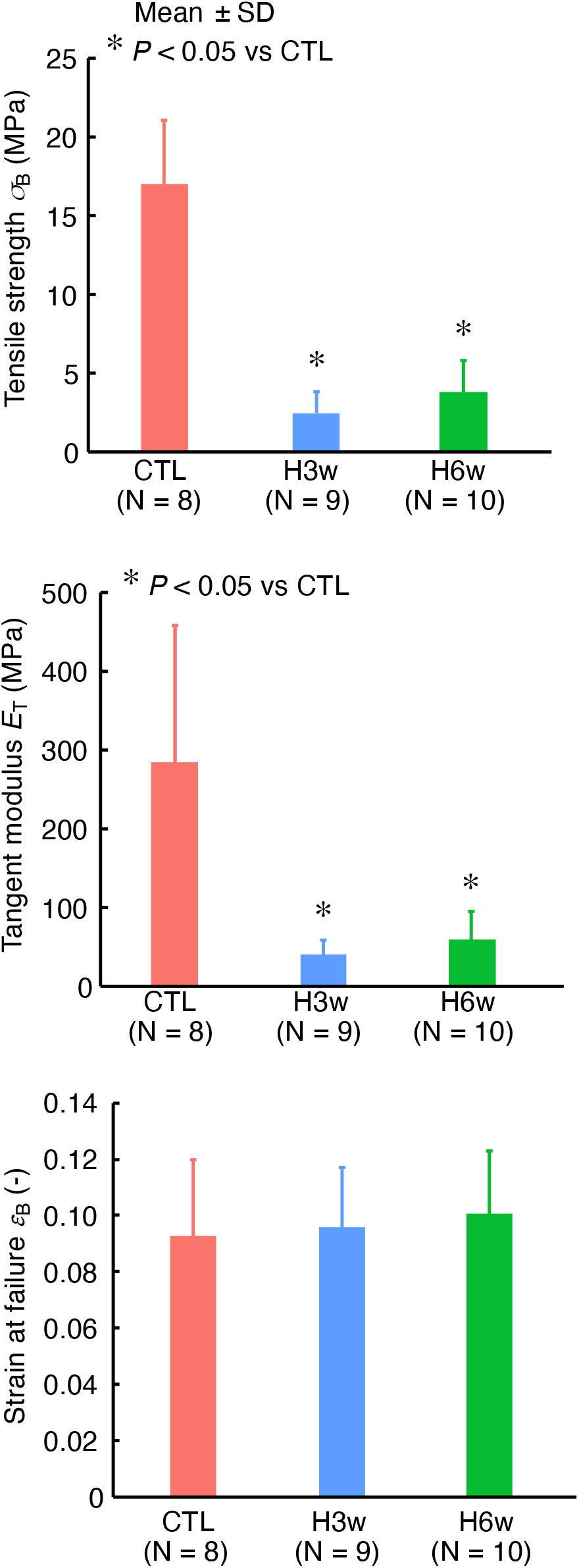
Tensile strength, tangent modulus, and strain at failure of mouse patellar tendons and their healing tissues. CTL, contralateral control PT; H3w, healing tissue regenerated in the PT defect examined at 3 weeks; and H6w, healing tissue regenerated in the PT defect examined at 6 weeks; N, number of specimens; SD, Standard deviation.

### 3.2 Collagen fibre alignment

The normal tendon exhibited collagen fibres aligned along the long axis of the tendon (Fig. 3(a)). Most of the cell nuclei in the normal PT resided in the interfiber spaces and were aligned along the collagen fibres. Conversely, H3w and H6w showed apparently different tissue structures from those in the normal tendon. At 3 weeks, collagen fibre alignment in the healing tissue was not as uniform as that in the normal tendon tissue. In addition, the number of cells was greater in the healing tissue than in the normal tendon, most of which possessed a round nucleus. The tissue structure and the number and shape of the cell nucleus at 6 weeks were similar to those at 3 weeks. Moreover, the distribution of the peak angle of collagen fibres in the analysed images demonstrated a high single peak at 0°, with a narrow distribution in the control tendon, and a lower peak with a broader distribution in the H3w and H6w (Fig. 3(b)).

**Figure 3.**
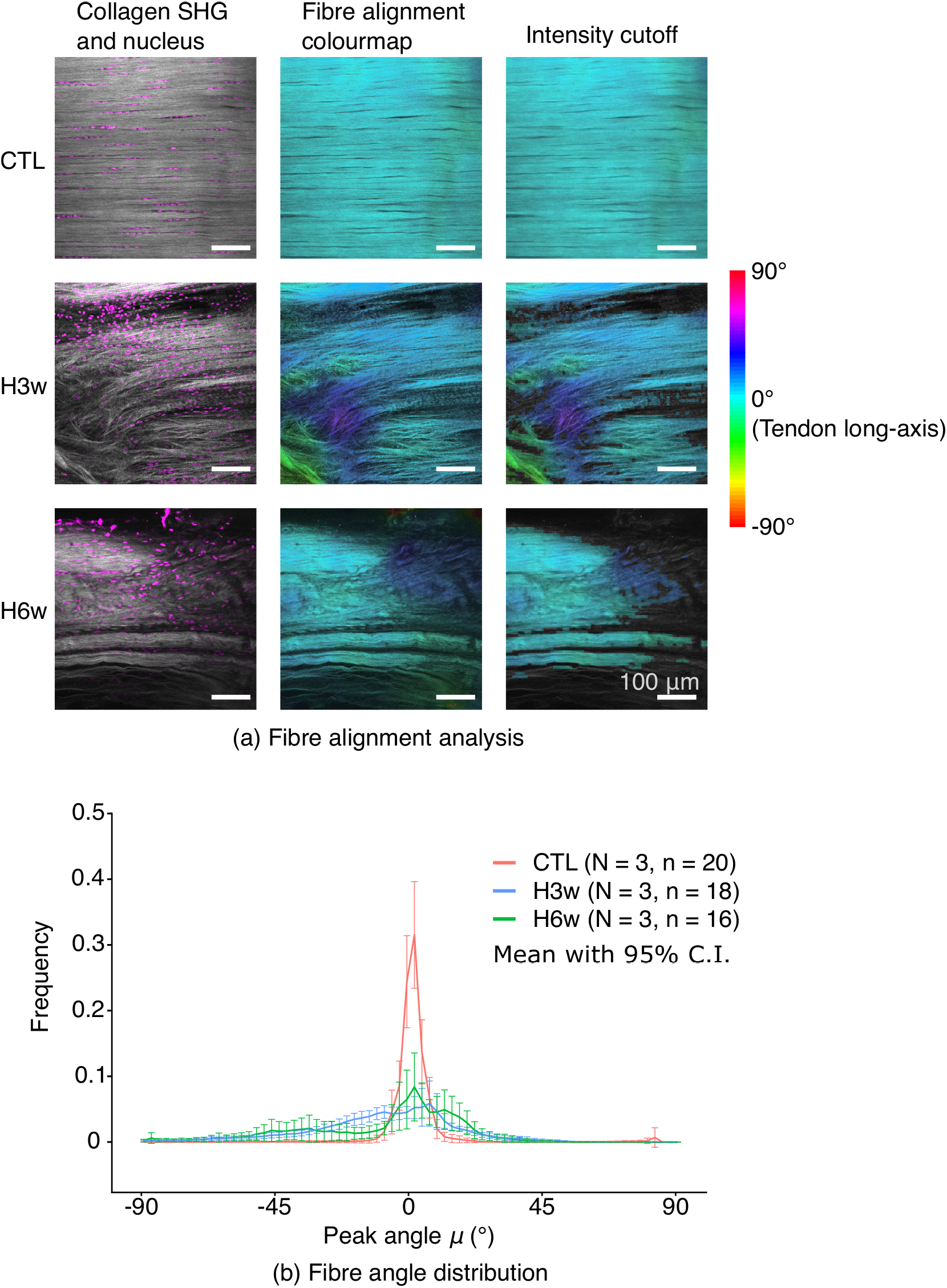
(a) Representative images from two-photon microscopic observation of collagen fibres and cell nuclei of CTL, H3w, and H6w; colourmap of collagen fibre alignment angle of these images and those following the intensity cut-off. (b) Distribution plot of collagen fibre alignment angle in CTL, H3w, and H6w. CI, confidence interval; N, number of specimens; n, number of images analysed; CTL, contralateral control PT; H3w, healing tissue regenerated in the PT defect examined at 3 weeks; and H6w, healing tissue regenerated in the PT defect examined at 6 weeks.

To quantitatively evaluate the alignment of collagen fibres, we obtained the orientation parameters *µ* (peak angle) and *κ* (distribution concentration) from the angular histogram of the alignment angle of collagen fibres from normal PT as well as H3w and H6w. It was demonstrated that the peak of the alignment angle distribution was almost 0 in all three groups but with large variability in the healing tissues. There was a statistically significant difference between the control and H3w group: 0.01° (median) in the control group compared with −0.11° (median) in H3w group (Fig. 4(a)). Furthermore, the distribution concentration was significantly higher in the control PT (median value = 475) than in H3w (8.85) and H6w (47.7).

**Figure 4.**
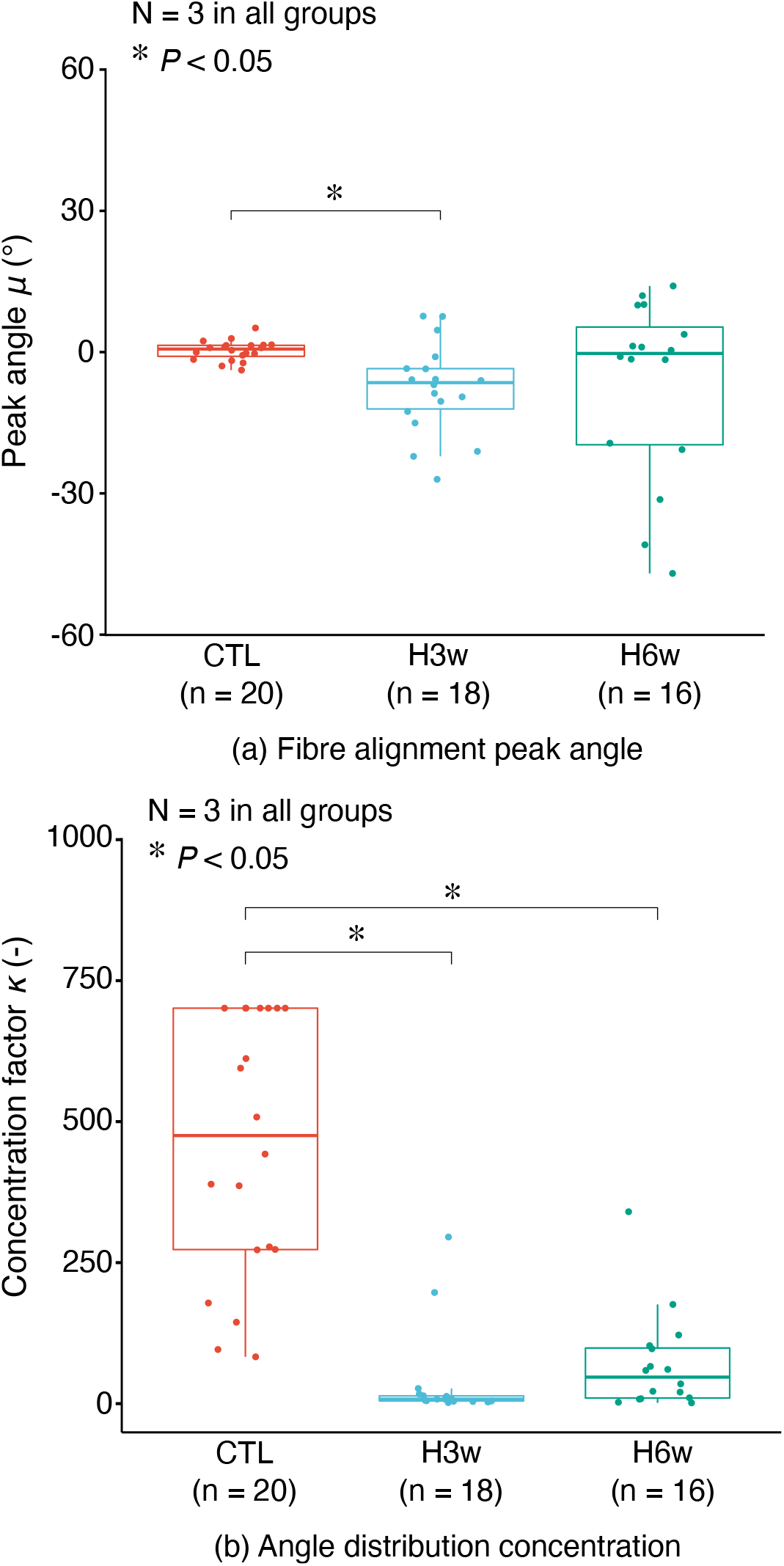
Peak angle *µ* (a) and concentration factor *κ* (b), as determined from collagen fibre alignment analysis. Box-and-whisker plots present median and IQR with the box and the maximum and minimum with whiskers. Small filled circles indicate individual data. Outliers located above the upper whisker and/or below the lower whisker were defined as values greater than the 75th percentile + 1.5IQR and/or values smaller than the 25th percentile – 1.5IQR. These outliers were also included in statistical analysis. N, number of specimens; n, number of images analysed; CTL, contralateral control PT; H3w, healing tissue regenerated in the PT defect examined at 3 weeks; and H6w, healing tissue regenerated in the PT defect examined at 6 weeks.

### 3.3 Microscopic strain analysis

Microscopic tissue strain was gradually increased with the application of the gross stretch in the normal tendon, reaching 4.2 ± 1.7% tensile strain at 700-µm gross stretch (Fig. 5(a)). In contrast, the strain increased at a greater rate with gross stretching in both H3w and H6w (7.9 ± 3.7% and 5.7 ± 3.3% tensile strain, respectively) at 700-µm gross stretch (Fig. 5(a)).

**Figure 5.**
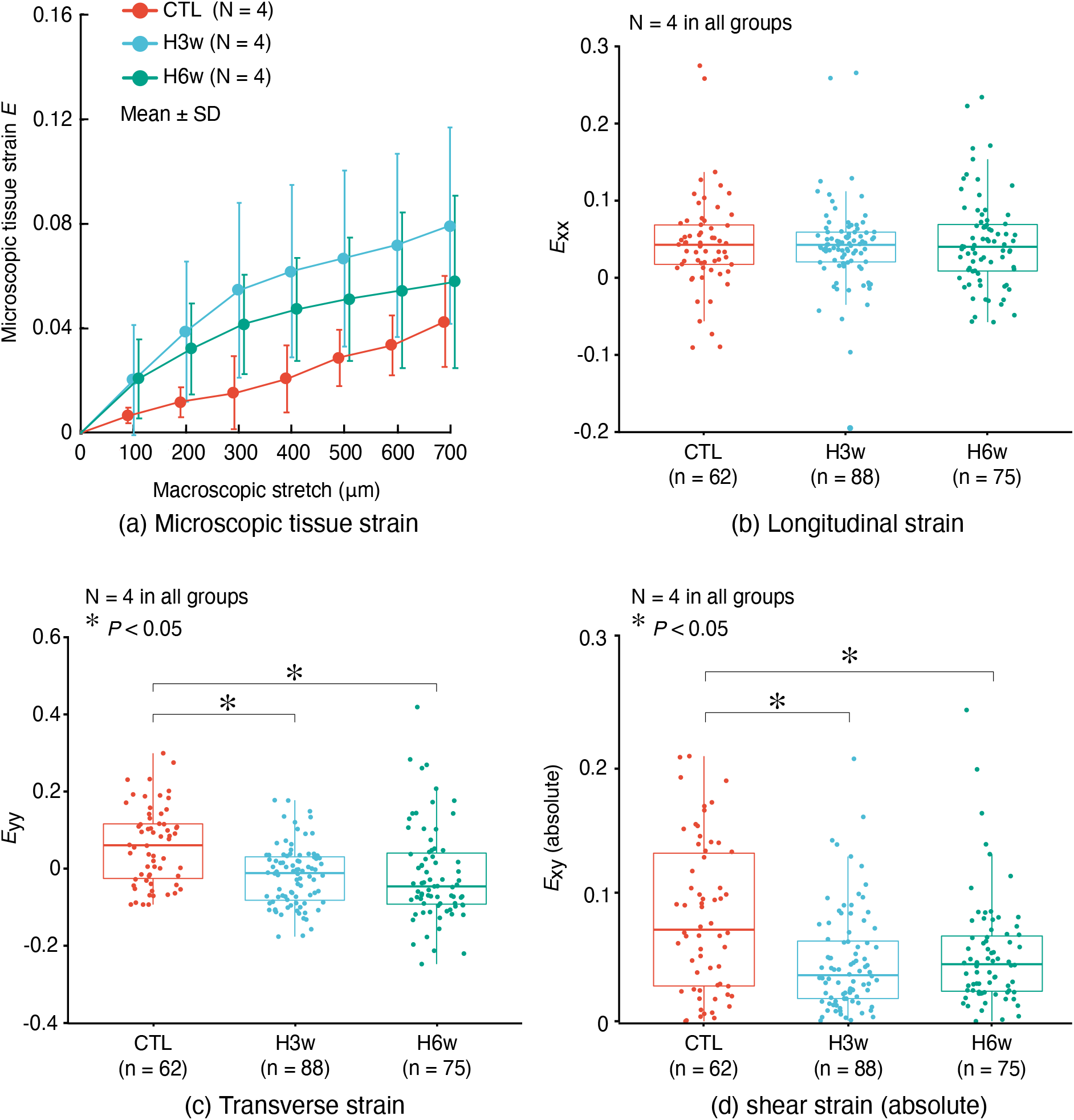
(a) Relationships between macroscopic stretch and microscopic tissue strain *E* in CTL, H3w, and H6w. (b-d) Longitudinal (*E*_xx_), transverse (*E*_yy_), and shear strain (*E*_xy_) at 4% microscopic tissue strain from Green-Lagrange strain tensor, calculated from the local strain field analysis. Box-and-whisker plots present median and IQR with the box and the maximum and minimum with whiskers. Small filled circles indicate individual data. Outliers located above the upper whisker and/or below the lower whisker were defined as values greater than the 75th percentile + 1.5IQR and/or values smaller than the 25th percentile – 1.5IQR. These outliers were also included in statistical analysis. N, number of specimens; n, number of elements analysed; CTL, contralateral control PT; H3w, healing tissue regenerated in the PT defect examined at 3 weeks; and H6w, healing tissue regenerated in the PT defect examined at 6 weeks.

To evaluate the microscopic local strain, Green-Lagrange strain tensor components *E*_xx_ (strain in the longitudinal direction), *E*_yy_ (strain in the transverse direction), and *E*_xy_ (shear strain) were evaluated in the three types of tissues at the 4% microscopic tissue strain level. Figure 6 shows a visual representation of the spatial variability of local strain values in each of the three directions in a representative specimen of the control, H3w, and H6w at 4% microscopic tissue strain level. In both normal and healing tissues, there were some local regions where greater strain values were calculated than those in other regions of the same specimen. Longitudinal strain *E*_xx_ exhibited no statistically significant differences among the three groups (Fig. 5(b)); the median values of *E*_xx_ were 0.042, 0.042, and 0.040 for the control, H3w, and H6w specimens, respectively. In contrast, the transverse strain *E*_yy_ in the control tendon (0.060) was significantly higher than that in H3w (−0.018) and H6w (−0.013). Moreover, the shear strain *E*_xy_ in the control tendon (0.080) was significantly higher than that in H3w (0.046) and H6w (0.044).

**Figure 6.**
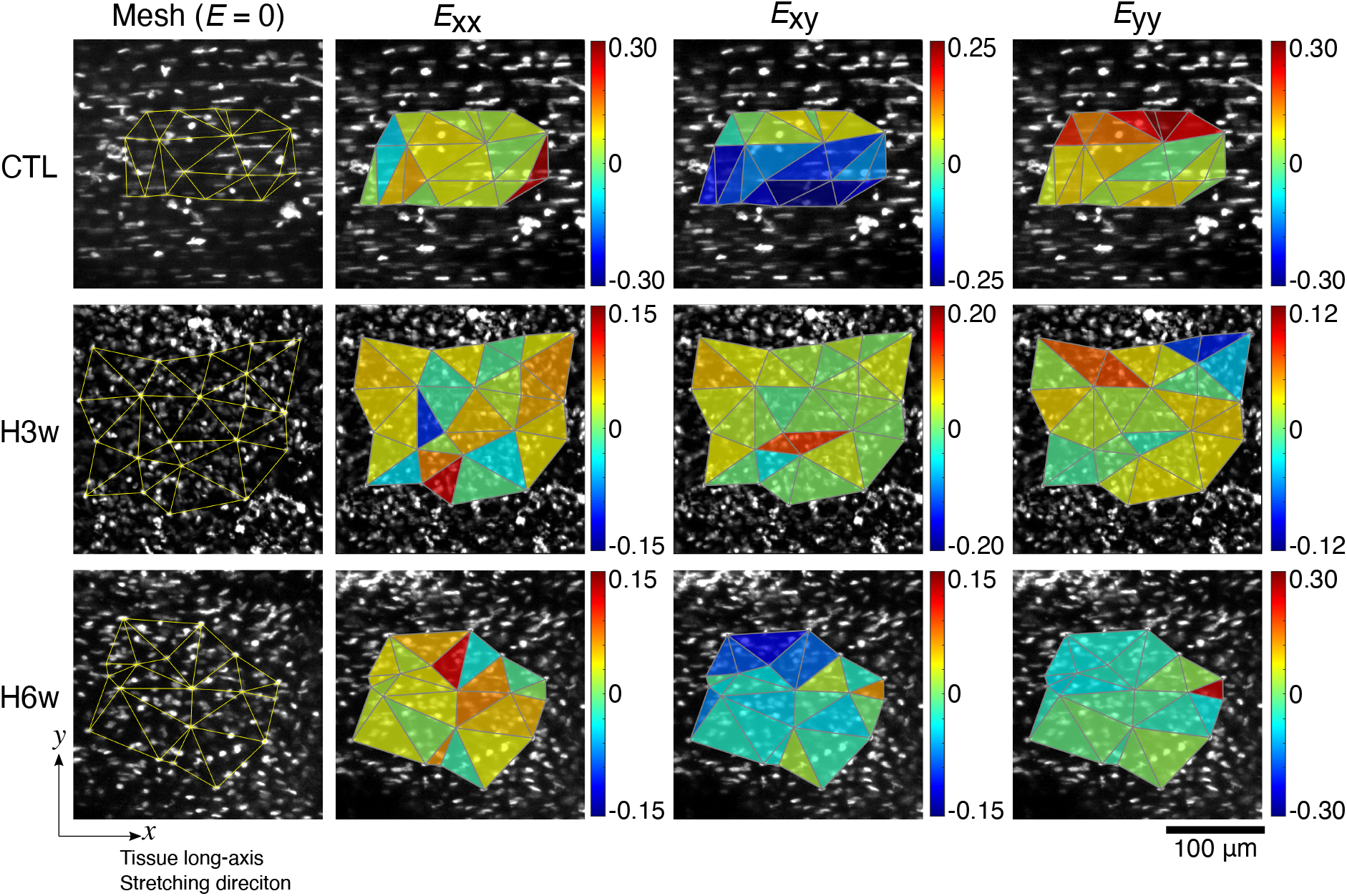
Spatial distribution of longitudinal (*E*_xx_), transverse (*E*_yy_), and shear strain (*E*_xy_) calculated in CTL, H3w, and H6w using triangular elements generated with the cell nuclei positions. Each specimen was stretched at 4% microscopic tissue strain (*E*). CTL, contralateral control PT; H3w, healing tissue regenerated in the PT defect examined at 3 weeks; and H6w, healing tissue regenerated in the PT defect examined at 6 weeks.

The cell nuclear aspect ratio (NAR) was 3.02, 1.88, and 1.77 at zero strain in CTL, H3w, and H6w, respectively, thus exhibiting statistically significant differences between control and H3w as well as CTL and H6w (Fig. 7(a)). *Δ*NAR was also relatively larger in CTL than that in H3w and H6w, the differences were not statistically significant (*P* = 0.054 for CTL vs H3w and 0.12 for CTL vs H6w, respectively).

**Figure 7.**
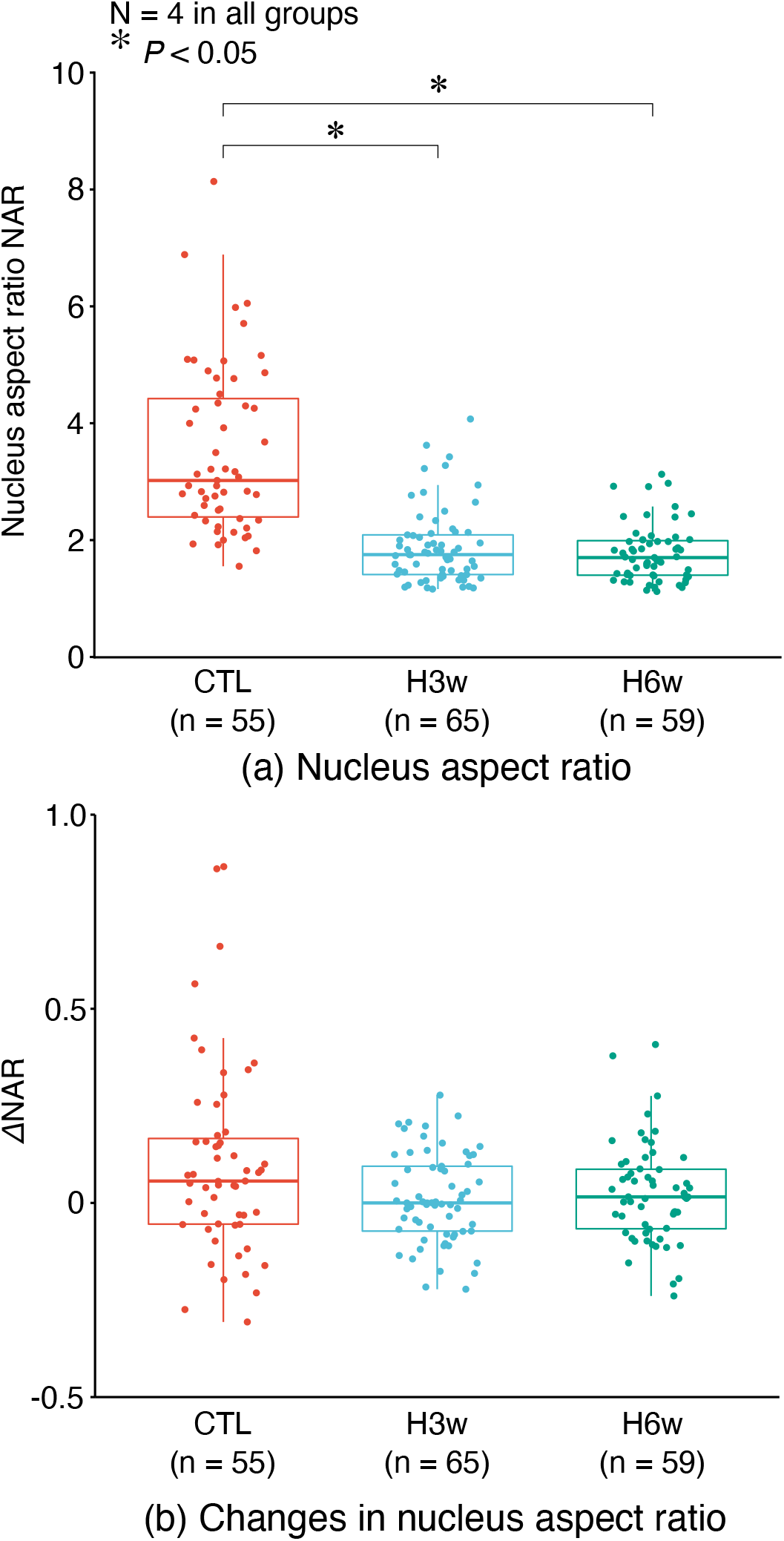
Cell nucleus aspect ratio at zero-strain level (a) and its change at 4% microscopic tissue strain (b) in CTL, H3w, and H6w. Box-and-whisker plots present the median and IQR with the box and the maximum and minimum with whiskers. Small filled circles indicate individual data. Outliers located above the upper whisker and/or below the lower whisker were defined as values greater than the 75th percentile + 1.5IQR and/or values smaller than the 25th percentile – 1.5IQR. These outliers were also included in statistical analysis. N, number of specimens; n, number of cells analysed; CTL, contralateral control PT; H3w, healing tissue regenerated in the PT defect examined at 3 weeks; and H6w, healing tissue regenerated in the PT defect examined at 6 weeks.

## 4. Discussion

The present study examined how the microscopic local mechanical environment differs between healthy, normal tendons and the healing tissue regenerated within a tendon defect, to better understand why tendon healing ends in scar formation. The results demonstrated that microscopic local strain remarkably differed between the normal tendon and healing tissues even if the microscopic tissue strain was at the same level. In particular, there was a significant difference in the amount of shear strain in the collagen fibre matrix, suggesting the importance of shear strain as mechanical stimulation to tenocytes for their proper functioning.

We adopted the central one-third defect model in PT, which has also been utilised in previous studies in the same animal species (Dyment et al., 2013, 2012) as well as in our previous studies using a rabbit model (Maeda et al., 2010, 2009, 2007). The gradual increase in the tensile strength and tangent modulus in healing tendon tissue observed in the present study was essentially similar to those observed in previous studies. It was also confirmed in our experiments that the strength and modulus of healing tissue at 9 weeks were 4.3 ± 1.4 and 83. 3 ± 39.0 MPa (25% and 29% of control PT), respectively, which were at the same level as that at 6 weeks. This finding is consistent with those of previous studies, showing that the mechanical properties of healing tissue following the first 5 or 6 weeks of healing did not improve greatly in the subsequent weeks; for example, in the rabbit PT central defect model, the strength of healing tissue was 18%, 25%, and 35% of control PT tissue at 3, 6, and 12 weeks, respectively (Maeda et al., 2009). Therefore, the investigation of the mechanical environment at 3 and 6 weeks of healing focused on characterising the healing tissue organisation and the amount of mechanical strain to which the cells within the tissue are exposed in the early phase of tendon defect healing. The low strength and elasticity of healing tissue may be attributed to differences in collagen matrix characteristics from normal tendons, such as fibre organisation (Figs. 3 and 4), as has also been reported previously (Freedman et al., 2018), and the types and amount of collagen molecules in the tissue (Sakabe et al., 2018; Williams et al., 1980). Although the types and amount of collagen were not quantified in the present study, it required a higher detector sensitivity setting in the two-photon imaging for collagen SHG signals in the healing tissues, indicating that the types and/or the amount of collagen within the healing tissues were qualitatively different from those of healthy, normal tendon tissue.

Such abnormalities of the collagen matrix in healing tissue also affect how fibres behave, and subsequently, how cells within the tissue are stimulated under mechanical loading. The behaviour of tendon structural components, collagen fibres and fascicles, have been well characterised in normal tendon tissue under uniaxial stretching (Cheng and Screen, 2007; Screen et al., 2004; Szczesny and Elliott, 2014; Thorpe et al., 2015, 2013). Tendon extension is achieved via straightening of collagen crimp morphology, followed by unwinding (rotation) of helical collagen fibres within fascicles, stretching of the collagen fibres, and sliding of adjacent fascicles and fibres. Accordingly, mechanical strains in the longitudinal, transverse, and shear directions are generated in the tendon collagen matrix. In the present study, in both normal and healing tissues, the collagen matrix was strained in the longitudinal, transverse, and shear directions, and the magnitude of transverse and shear strains were significantly different between normal and healing tissues. For the difference in the magnitude of the transverse strain, the positive transverse strain calculated in normal tendon may be attributable to the three-dimensional rotation and realignment of collagen fibre bundles or collagen fibres themselves. Thus, it is likely that the fibre matrix was not actually expanded, and cells within the matrix were subjected to tensile strain in the transverse direction while the tendon itself was longitudinally stretched macroscopically. Accordingly, it may be difficult to evaluate the actual difference in the magnitude of the matrix strain in the transverse direction based on the data presented here. For shear strain, despite a large amount of variability observed in both normal and healing tissues, the magnitude of shear strain in the normal tendon was significantly greater than that in both H3w and H6w. Because the magnitude of longitudinal strain and that of shear strain were not correlated in the control tendon (Fig. 8) but a greater magnitude of shear strain than longitudinal strain was evident in 70% of the triangular elements analysed, sliding between collagen fibres was already evident at this level of tissue strain, as has also been reported in previous studies (Screen et al., 2004; Szczesny and Elliott, 2014; Thorpe et al., 2015, 2013).

**Figure 8.**
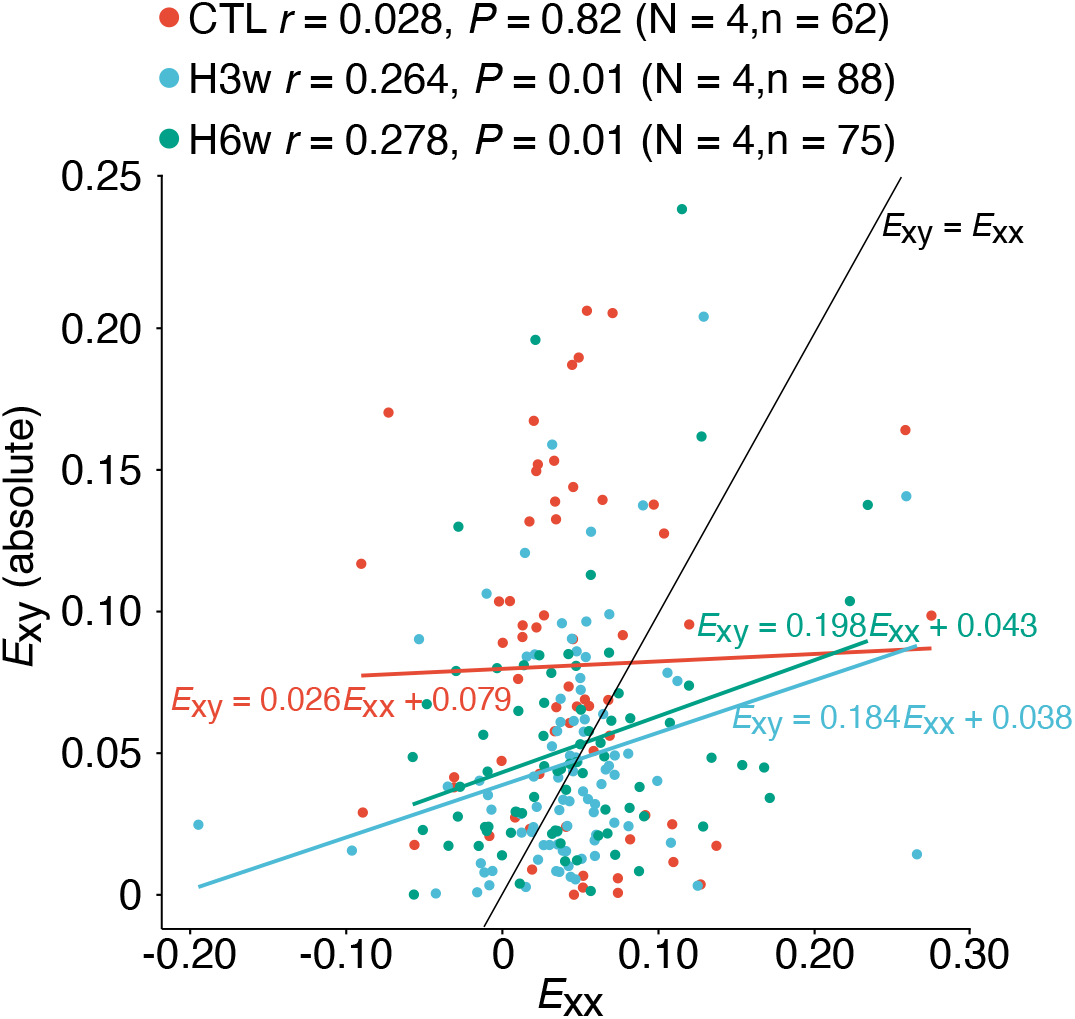
Correlation analysis between local longitudinal strain *E*_xx_ and local shear strain *E*_xy_ in CTL, H3w, and H6w at 4% microscopic tissue strain. *r*, Pearson’s product moment correlation coefficient; *P*, p-value from the correlation analysis; N, number of specimens; n, number of elements; CTL, contralateral control PT; H3w, healing tissue regenerated in the PT defect examined at 3 weeks; and H6w, healing tissue regenerated in the PT defect examined at 6 weeks.

In contrast, there was a weak but significant correlation between shear and longitudinal strains in both H3w and H6w. It was shown that 49% and 44% of the triangular elements had a lower magnitude of shear strain than that of longitudinal strain. This may imply that collagen fibre sliding was not evident in stretched healing tissue, possibly due to that healing tissue may have no clear hierarchical structure. The differences in collagen fibre behaviour during macroscopic stretching between normal and healing tendons have also been characterised from the view point of the degree of collagen fibre disorganisation (Freedman et al., 2018). In this study, it was demonstrated that collagen fibre disorganisation was greater in healing tissue than in uninjured, normal tendon tissue, and the strain-induced change of collagen fibre alignment observed in the normal tendon was less effective in healing tissue. Accordingly, because healing tendon tissue in the early phase of healing is less organised in the longitudinal direction and does not form the hierarchical structure as that in the normal tendon, collagen fibre realignment, rotation, and sliding are not induced during macroscopic stretching.

These differences in the strain field between normal tendon and healing tissues were also reflected in the shape of cells within the tissues, i.e. the cells in the normal tendon had a more elongated shape (higher aspect ratio) than those in healing tissues. A large magnitude of shear deformation (sliding between collagen fibre bundles) is possibly transferred to the tenocytes residing between the bundles of collagen fibres and thus contributes to cell deformation along with the longitudinal strain, resulting in an elongated shape of the cell and its nucleus. Conversely, a small magnitude of shear strain in healing tissues may not contribute to the cell deformation as much as that in case of normal tendons, resulting in a less elongated shape. Although there were no significant differences in the change in nuclear aspect ratio between normal and healing tendons (Fig. 8(b)), the change was relatively larger in CTL, suggesting that the cells in healing tendon tissues may experience lower mechanical stress than that experienced by cells in normal tendon tissues, which is consistent with previous findings (Freedman et al., 2018).

It is well known that the application of mechanical stress in tendons is an essential factor in the proper functioning of tenocytes and therefore in the maintenance of tissue integrity of tendons (Wang, 2006). In addition, the presence of mechanical stress is necessary for better tendon healing because the deprivation of mechanical stress from the healing tendon results in a markedly lower strength and elastic modulus in healing tendon tissues (Maeda et al., 2009). Nonetheless, the strain field within the healing tendon tissue was different from that within the normal tendon tissue, lacking shear strain derived from sliding between fibres and fibre bundles. Therefore, although some mechanical stress is applied to the cells in healing tendon tissues, a better mechanical environment may be needed to appropriately stimulate the cells in the healing tissues. This could be achieved by the introduction of a tissue-engineered scaffold with a highly aligned fibre structure, recapitulating the dynamics of native tendon collagen fibres, in which cells penetrating from the remaining tendon tissue as well as surrounding tissues are stimulated as they were within the normal tendon matrix.

It has been demonstrated that the repair process of the central defect of the PT is predominantly governed by tendon progenitor cells from paratenon, a sheath-like tissue encompassing the whole tendon; the multipotent cells migrate into the defect, become tenogenic by expressing tendon-specific transcription factor scleraxis, and produce extracellular matrix components, such as type III collagen at an early stage of healing to fill the defect (Dyment et al., 2013). The expression of scleraxis is essential for the success of the remodelling of type III collagen matrix of the healing tissue to a more tendon-like matrix, consisting of type I collagen and other associated molecules (Sakabe et al., 2018). Moreover, the expression of scleraxis in tendon cells is responsive to mechanical loading *in vivo*, and an *in vitro* experiment has shown that the application of mechanical stimulation, in particular fluid shear stress, is effective in the induction of scleraxis expression in tenocytes (Maeda et al., 2011). These findings may highlight the importance of mechanical stimulation to healing tissue cells by shear deformation for better tissue repair.

It was one of the limitations in the present study that we analysed the microscopic strain at only one deformation point (4% microscopic tissue strain level); this was because the microscopic tissue strain in the control tendon was markedly smaller than that in H3w and H6w at each 100-µm macroscopic stretch step (Fig. 5(a)). One possible reason for this is that the stiffness of the control PT was greater than that of the custom-made acrylic grips, and thus, the grips were also deformed during PT stretching. The second possible reason is that the deformation mainly occurred at the attachment site to the patella and/or tibia, so that the amount of deformation was attenuated at the mid-portion of the tissues. On the other hand, the remaining tendon tissue at both sides of the central defect was possibly weakened and became more compliant with the increase in the mechanical load applied owing to the decrease in tendon cross-sectional area by creating the defect (Maeda et al., 2009). These differences in macroscopic behaviour between the normal tendon and healing tissue only allowed the comparison of microscopic strain field to be made where microscopic tissue strain was at the same level between the two types of tissues.

Another limitation that should be mentioned is the use of small triangular elements in microscopic local strain analysis. This methodology has previously been utilised and verified in normal tendon tissues (Han et al., 2013; Screen and Evans, 2009) but not in healing tissues with a less-organised fibre structure. To avoid the objective generation of the elements, the Delaunay method was used, and elements with skewed shapes (thin and long triangles) were manually eliminated. However, although collagen fibre alignment could be easily speculated from the locations of the cell nuclei, and therefore, the three cell nuclei of each element were structurally relevant (a triangle was formed across the same bundle of collagen fibres), which might not be the case in healing tissues. Thus, triangular elements in healing tissues might have been generated with three cell nuclei with low structural relevance. This could have resulted in a large variability in calculated strain values compared to that in the normal tendon; however, the variability of strain values was at almost the same values (interquartile range was 0.051, 0.029, and 0.060 for *E*_xx_, 0.141, 0.112, and 0.132 for *E*_yy_, and 0.103, 0.044, and 0.042 for *E*_xy_(abs) in control, H3w, and H6w, respectively); therefore, the strain analysis performed in this study is considered acceptable. Nonetheless, in future studies, strain analysis in healing tissues using the cell nuclei should be performed with the visualisation of fibre structure with multiphoton imaging or fluorescent staining of matrix fibres.

## 5. Conclusion

In conclusion, the present study demonstrated that microscopic tissue deformation was significantly different between normal tendon and healing tendon tissues at the early healing phase, in particular a significantly smaller magnitude of shear strain in healing tissue than normal tendon under uniaxial stretch. This difference is supposed to be reflected in the difference in cell nuclear shape and possibly the amount of mechanical stimuli applied to the cells during tendon deformation. Accordingly, restoration of a normal local mechanical environment in the healing tissue may be key to a better healing outcome of tendon injury.

## Acknowledgements

The present study was supported in part by Japanese Society of Promotion Science KAKENHI grant numbers 18H03752 (TM), 19K22960 (TM) and 20K21887 (EM). The authors wish to acknowledge Division for Medical Research Engineering, Nagoya University Graduate School of Medicine, for the use of two-photon microscope (A1RMP, Nikon). We also appreciate greatly Professor Nobuto Kitamura at Department of Orthopaedic Surgery, St. Luke’s International Hospital, Tokyo, Japan, for helpful discussion.

